# Host-initiated microbial association leads to stable ectosymbiosis in an ecological model

**DOI:** 10.1101/2024.09.08.611858

**Authors:** Nandakishor Krishnan, István Zachar, Ádám Kun, Chaitanya S. Gokhale, József Garay

## Abstract

Microbial symbiosis is extremely common among metabolically coupled cells, and, presumably, gave rise to mitochondria. How such symbioses emerge, evolve and stabilize are unknown, especially in the prokaryotic domain, where endosymbioses are virtually nonexistent. Yet there is growing evidence suggesting that the eukaryotic cell emerged from such a prokaryotic partnership, where integration was not the result of phagocytotic inclusion (as in case of plastids) but of metabolic cooperation. While prokaryotes almost ubiquitously engage in metabolic syntrophy, it is unknown if such cooperation alone can enable stable, dependent ectosymbioses that could pave the road toward physical integration of parties. Here, we tested the hypothesis that free-living syntrophy can lead to stable ectosymbiosis between microbial partners, using an ecological mathematical model. Assuming an already syntrophic and asymmetric partnership of free-living hosts and symbionts, we investigated under what conditions obligate ectosymbiosis evolve. Our results show that reduced inhibition (of self-inhibiting metabolic products) over the contact surface of partners can stabilize the ectosymbiotic consortia against free-living forms. Furthermore, strong metabolic activity between the host and their ectosymbionts could facilitate obligacy in their physical association. The model points to the significance of the contact surface in the evolution of crucial endosymbiotic features. Our results support the hypothesis that cooperative, syntrophic microbes (especially prokaryotes) are capable of coevolving to form species-specific ectosymbiosis by means of reducing the inhibition of accumulating products, a necessary first step towards endosymbiotic integration. The model provides a plausible explanation on how common metabolic syntrophy can lead to physical integration of parties through gradual ectosymbiosis. Moreover, our work fills a gap between microbial cooperation models (assuming free-living species) and those that are concerned only with already concluded physical integration under a multilevel selection paradigm.

## Introduction

Microbes, especially prokaryotes, almost ubiquitously engage in metabolic cooperation at every habitat they occupy. They establish cooperation through various metabolites they produce or enable in their neighborhood via their metabolisms. They can evolve dependencies on biofunctions of each other on a diverse scale, ranging from pairwise to multipartite associations (Krause et al., 2017), and from direct cross-feeding to protection mutualism (D’Souza et al., 2018; Smith et al., 2019; Zachar and Boza, 2022). Syntrophy is an umbrella term to denote metabolic interaction in which constituent partners rely on one another (bi- or unilateral) for metabolites for their growth (McInerney et al., 2008; Morris et al., 2013; Schink, 2002, 1997; Sieber et al., 2012). The microbial syntrophic relationship facilitates a synergy for associations and communities among them (Morris et al., 2013; Rothschild and Mancinelli, 2001; Shu and Huang, 2022; van Wolferen et al., 2018). Permanent and obligate pairwise metabolic cooperation may even evolve into physically linked symbioses, where one partner is the long-term ectosymbiont of the other (*Chlorochromatium* (Cerqueda-García et al., 2014; Liu et al., 2013; Overmann and van Gemerden, 2000) or epibiotic GPB on *Giganthauma karukerense* Thaumarchaeota (Muller et al., 2010)). When a syntrophic partnership ensures better-than-random probability of co-inheritance of partners, group selection may select for the pair instead of free-living individuals. As a result, a new evolutionary unit may evolve under a new level of selection, ultimately leading to a major evolutionary transition (Maynard Smith and Szathmáry, 1995; Szathmáry, 2015). Importantly, this need not happen when one partner is physically integrated into the other (even in case of ectosymbiosis), but when they become obligately dependent on each other.

According to mainstream hypotheses (Imachi et al., 2020; López-García et al., 2017; López-García and Moreira, 2023; Martin et al., 2015; Spang et al., 2015), the origin of the eukaryotic cell, a major transition itself, was the result of such a metabolic cooperation between free-living cells that turned obligate and irreversible, one partner ultimately ending up within the other. While the ancestral partners are pointed out with increasing confidence from the bacterial and archaeal domains (Eme et al., 2017; Imachi et al., 2020; López-García and Moreira, 2023; Muñoz-Gómez et al., 2022; Spang et al., 2015), there is no consensus on many critical points of the origin (Richards et al., 2024). Their initial interaction, benefits and mechanism of inclusion are still unknown, and the omics data trickling in can rarely differentiate among contending hypotheses, leaving room for much speculation, often without merit (most, if not all, hypotheses remain to be validated (Zachar and Szathmáry, 2017)). The issue is further complicated by the fact that eukaryogenesis happened more than a billion years ago with elusive traces, and without second examples. There is no other origin of the nucleus and no other merger between two prokaryotes being on par with the success and stability of the ancestral mitochondrion and its host. Despite a few known cases of ectosymbiotic prokaryotic partnerships (Dombrowski et al., 2019; Zachar and Boza, 2020), we do not know how these partnerships were initiated, stabilized and coevolved. As a result, it remains unclear how obligate symbioses of syntrophic partners may ensue. Without directly observable cases of prokaryotic ecto- or endosymbioses emerging from metabolic cooperation (natural or artificial), theoretical modelling remains the only approach to identify the potential driving factors that can lead to stable symbiotic consortia in a syntrophically interacting community. Yet the field of eukaryogenesis severely lacks appropriate modelling to back up the various theories, and research has notoriously been hampered by the lack of models to test the hypotheses.

One particularly long-lasting debate concerns the original interaction of eukaryogenetic parties, specifically the nature and formation of the initial interaction of the host and the future mitochondrion. Arguably, how the symbiont got inside the host is extremely important. While phagotrophy or parasitic invasion cannot yet be excluded, here we focus on metabolic syntrophy only. More specifically, we focus on a step that likely preceded physical integration. We propose that a stable ectosymbiosis evolved out of a syntrophic system before the internalization of the symbiont (also see Baum and Baum (2014)), contrary to the possibility of direct internalization as in, e.g., phagotrophic farming (Zachar et al., 2018). While the metabolic coupling of the ancestral parties was and still is the focus of considerable research (López-García and Moreira, 2023, 2020; López-García and Moreira, 1999; Martin and Müller, 1998; Moreira and López-García, 1998; Searcy, 2003; Zachar and Szathmáry, 2017), syntrophic theories agree on one aspect: the metabolite-mediated cooperation between parties provided the selective advantage for the pair over free-living individuals. It is well established through modelling that metabolic cooperation can lead to microbial mutualism (Boza et al., 2023; Bull and Harcombe, 2009; Doebeli, 2002; Estrela et al., 2012; Estrela and Gudelj, 2010; Krishnan et al., 2024; Libby et al., 2019; Liu and Sumpter, 2017; Preussger et al., 2020), with models covering many specific metabolic interaction types (also see D’Souza et al. (2018) and Kost et al. (2023)). Yet, to our knowledge, there are no models that study how metabolic cooperation (syntrophy) may lead to stable, obligate physical integration, something that seems to be an absolutely crucial step on the road to endosymbiosis. The lack of such models or data urges us to investigate the hypothesis of syntrophy leading to endosymbiosis. Our model is motivated by the various syntrophic microbial cooperation cases known, with a focus on eukaryogenesis. We concentrate on a mechanism to bridge microbial models that assume free-living species in a community (even if part of biofilms (Hansen et al., 2007)) with those that deal with upfront physical integration (von der Dunk et al., 2022; Zachar et al., 2018).

Here, we introduce a theoretical eco-evolutionary model to test the specific hypothesis that metabolic syntrophy can lead to physical ectosymbiosis under a multilevel selection scenario. We assume a worst-case scenario of a physically independent (free-living) pair of *hosts* and *symbionts* belonging to different species in unidirectional syntrophy. The symbiont feeds on the metabolic product of the host. Excess concentration of the hosts’ metabolite (accumulated in the habitat by whole host population) is toxic to its producer, inhibiting growth. There are many examples among microbes where products inhibit growth and consumption (effectively reducing the concentration in the vicinity) by other species provides cooperative help, e.g. *Pseudomonas* sp. produces methanol that inhibits its growth while *Hyphomicrobium* sp. can utilize methanol (Morris et al., 2013; Wilkinson et al., 1974). Arguably, cases of metabolic interactions affecting microbial consortia are also prevalent (Liu et al., 2019; Piccardi et al., 2019; Wu et al., 2020).

Our model considers the invasion of a (costly) mutant host that can bind its free-living symbiont partners on its cell surface, forming ectosymbiotic association. In the ectosymbiotic consortia, symbionts on the host surface effectively reduce the free surface area of the host’s plasma membrane. This has two direct effects on the host: reduced surface implies constrained resource consumption, and it reduces exposure to the inhibitory effect of its (and conspecifics’) metabolic product. Most autotrophic microbes (especially prokaryotes) perform bioenergetics through their plasma membranes, which means that their bioenergetic production is directly dependent on cell surface area over which essential resources are taken up (Lane and Martin, 2012). Having partners attached to the surface imposes a severe trade-off, and it is not trivial when it is beneficial to pay the cost of reduced consumption despite reduced self-inhibition. Accordingly, we consider that the consortial host’s external surface area unoccupied by ectosymbionts is (and could have been) a significant factor in the formation of the stable ectosymbiotic consortia, possibly obligate as well.

## Results

We tested our hypothesis within a model where metabolically cooperating species may form physical associations, increasing the contact-surface of partners. We were interested in whether a mutant host, with the capability to capture or bind symbionts on its external cell surface, can invade the stable resident population of free-living host and symbiont species. Our methodology relies on density-dependent ecological (two-species) dynamics determining mutant invasion and fixation, aided by metabolite concentration dynamics. Under this assumption, we checked whether physical coupling is selected for, given the possibility of the host species to associate with free-living symbionts. Alternatively, whether such association becomes a hindrance due to the increased cost realized in reduced consumption rate for the mutant host. If the host is better off with increased consumption and baseline metabolic partnership, we expect that physical coupling will not fix in the population and metabolically dependent species continue their free-living lifestyle (as extant syntrophic prokaryotes do).

We model the evolutionary scenario leading to the formation of consortia from physically independent (free-living) individuals of two distinct species in three steps. We start with a system with a single species (host), enduring self-inhibition of growth by its metabolic product (Figure 1A; Appendix A1). Next, we consider a system with a new species (symbiont), enjoying benefits of an obligate syntrophic relationship with the resident host (Figure 1B; Appendix A2). A stable ecological coexistence of the two free-living species in this system guarantees the maintenance of syntrophy between them. Finally, we check whether a mutant host capable of initiating ectosymbiosis with the free-living symbionts (forming a two-species ectosymbiotic consortium) can invade the syntrophic resident system of free-living individuals (Figure 1C; Appendix A3). The population newly introduced to the system adds a dimension to the existing resident system, extending the ecological dynamics. The outcomes of ecological selection are based on the stability of the steady states of the resident-mutant dynamical system (Cressman, 1992; Cressman et al., 2020; Cressman and Garay, 2003a, 2003b). Eventually, we focus on the fixation of the ectosymbiotic consortia in the system based on the stability of the steady state with viable mutant host population density. The conditions for the *evolutionary substitution* (Cressman et al., 2020) of the resident host by its mutant phenotype, reveal the potential driving factors behind the fixation of the stable symbiotic consortia. Figure 1 is a schematic representation of the ecosystems we consider. For details of the individual systems, see Model and Methods and the corresponding Appendices.

**Figure 1.**
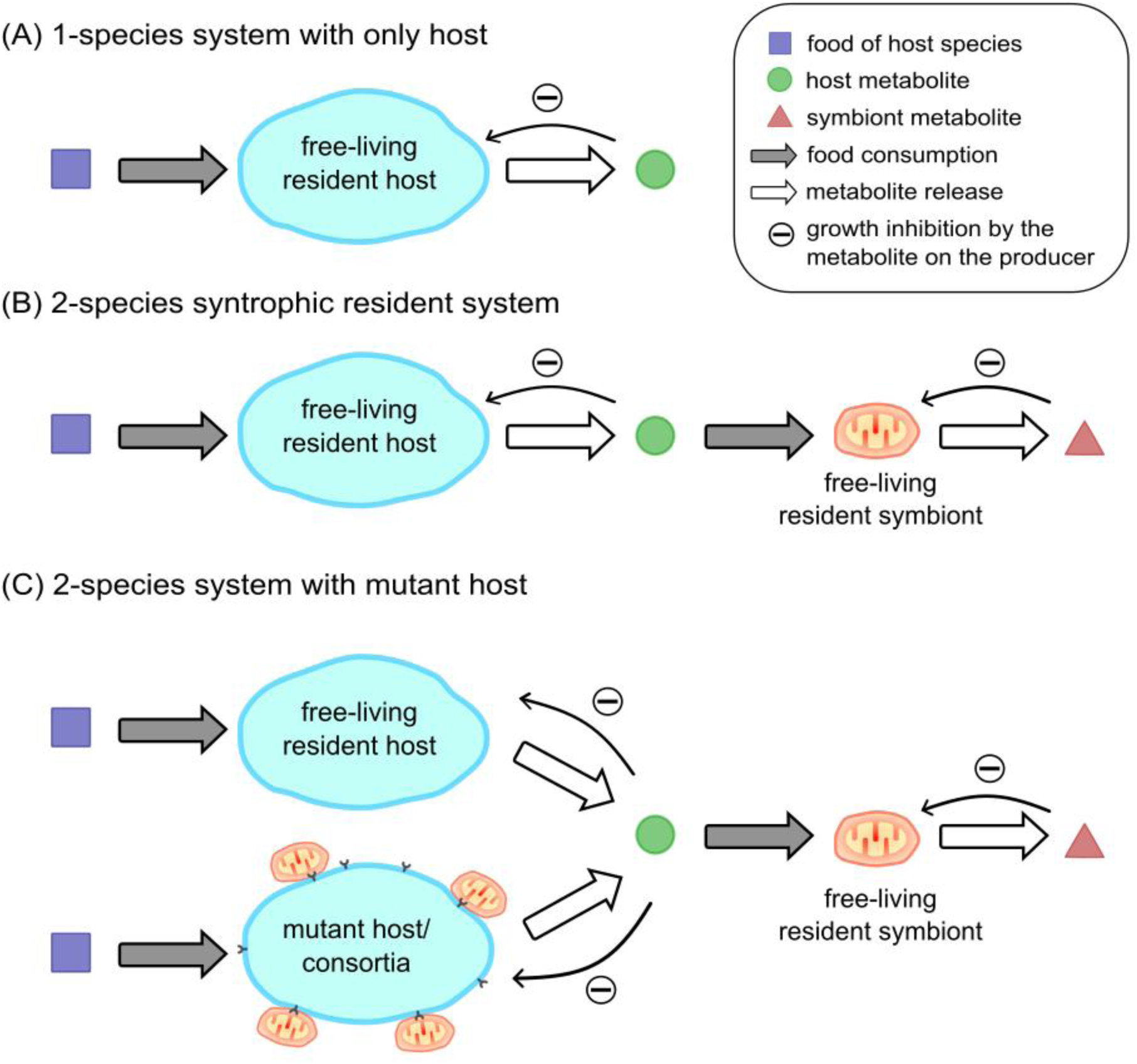
The three evolutionary steps toward ectosymbiotic consortium-formation. (A) Initially, there is a single host species producing a metabolite that exerts self-inhibition. (B) When the symbiont species is added to the system, it alleviates the effect on the host by consuming its self-inhibiting product, establishing unilateral syntrophy. The symbiont in turn produces a self-inhibiting metabolite too. (C) Mutant hosts capable of forming consortia may appear and invade the population of symbiotic syntrophs, leading to explicit ectosymbiosis.

The one-species resident dynamical system is analyzed for the stability of the unique interior fixed point, ensuring limited growth and stabilization of the host (species *X*) population density. The unique interior fixed point exists if and only if the growth rate of species *X* is less than the maximal growth inhibition (otherwise, the population explodes). The interior fixed point, if it exists, is unconditionally stable (see Figure 2; for details, see Appendix A1). This proves that metabolic growth inhibition is sufficient to prevent population explosion and stabilize population growth.

**Figure 2.**
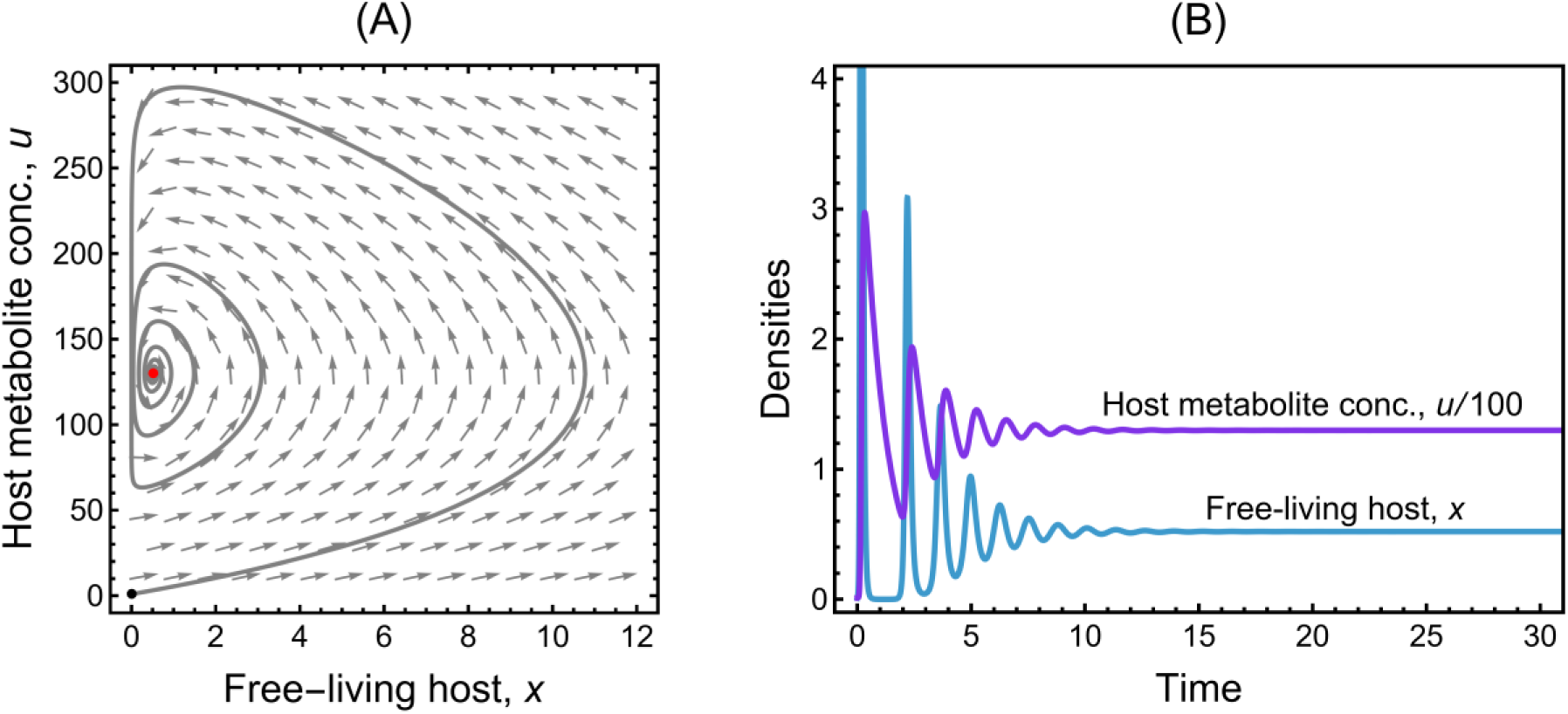
Stability of a self-inhibiting host species. (A) Phase portrait of the one-species system with the trajectory starting from an arbitrary point in the neighborhood of the trivial fixed point (black point). The dynamics stabilizes at the unique interior fixed point, (*x*^∗^, *u*^∗^) = (0.52, 130) (red point). (B) Solution of the one-species system dynamics for perturbed (0, 0) as the initial condition. The trivial fixed point (0, 0) is always unstable. The parameter values are as in Table 1. For details of the stability analysis, see Appendix A1.

**Table 1:**
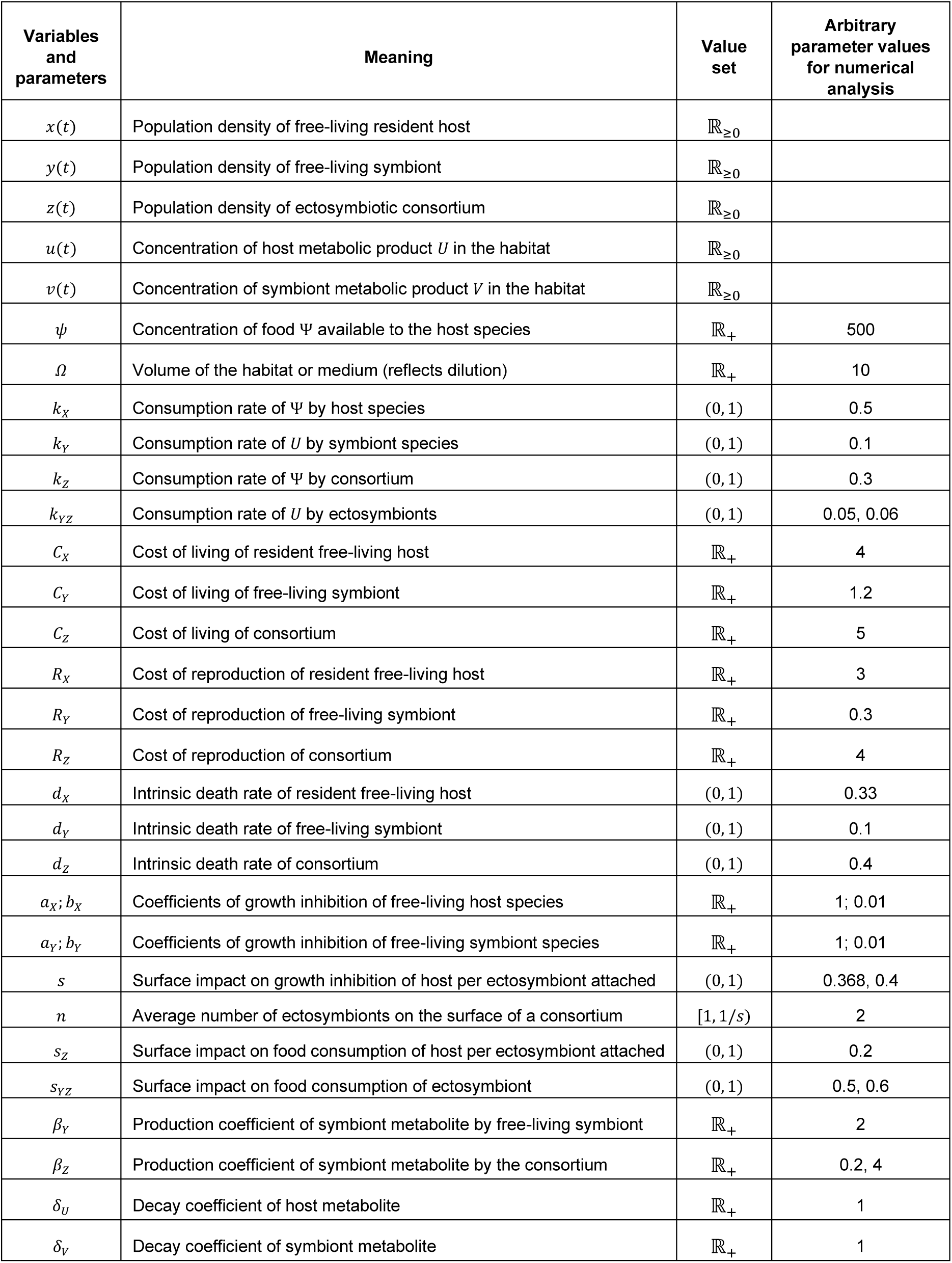
Variables and parameters of the model. . The parameter values are qualitatively consistent with the assumptions in the model descriptions and are used only to visualize analytical results.

| Variables and parameters | Meaning | Value set | Arbitrary parameter values for numerical analysis |
| --- | --- | --- | --- |
| $x(t)$ | Population density of free-living resident host | $\mathbb{R}_{\geq 0}$ | |
| $y(t)$ | Population density of free-living symbiont | $\mathbb{R}_{\geq 0}$ | |
| $z(t)$ | Population density of ectosymbiotic consortium | $\mathbb{R}_{\geq 0}$ | |
| $u(t)$ | Concentration of host metabolic product $U$ in the habitat | $\mathbb{R}_{\geq 0}$ | |
| $v(t)$ | Concentration of symbiont metabolic product $V$ in the habitat | $\mathbb{R}_{\geq 0}$ | |
| $\psi$ | Concentration of food $\Psi$ available to the host species | $\mathbb{R}_+$ | 500 |
| $\Omega$ | Volume of the habitat or medium (reflects dilution) | $\mathbb{R}_+$ | 10 |
| $k_X$ | Consumption rate of $\Psi$ by host species | $(0, 1)$ | 0.5 |
| $k_Y$ | Consumption rate of $U$ by symbiont species | $(0, 1)$ | 0.1 |
| $k_Z$ | Consumption rate of $\Psi$ by consortium | $(0, 1)$ | 0.3 |
| $k_{YZ}$ | Consumption rate of $U$ by ectosymbionts | $(0, 1)$ | 0.05, 0.06 |
| $C_X$ | Cost of living of resident free-living host | $\mathbb{R}_+$ | 4 |
| $C_Y$ | Cost of living of free-living symbiont | $\mathbb{R}_+$ | 1.2 |
| $C_Z$ | Cost of living of consortium | $\mathbb{R}_+$ | 5 |
| $R_X$ | Cost of reproduction of resident free-living host | $\mathbb{R}_+$ | 3 |
| $R_Y$ | Cost of reproduction of free-living symbiont | $\mathbb{R}_+$ | 0.3 |
| $R_Z$ | Cost of reproduction of consortium | $\mathbb{R}_+$ | 4 |
| $d_X$ | Intrinsic death rate of resident free-living host | $(0, 1)$ | 0.33 |
| $d_Y$ | Intrinsic death rate of free-living symbiont | $(0, 1)$ | 0.1 |
| $d_Z$ | Intrinsic death rate of consortium | $(0, 1)$ | 0.4 |
| $a_X; b_X$ | Coefficients of growth inhibition of free-living host species | $\mathbb{R}_+$ | 1; 0.01 |
| $a_Y; b_Y$ | Coefficients of growth inhibition of free-living symbiont species | $\mathbb{R}_+$ | 1; 0.01 |
| $s$ | Surface impact on growth inhibition of host per ectosymbiont attached | $(0, 1)$ | 0.368, 0.4 |
| $n$ | Average number of ectosymbionts on the surface of a consortium | $[1, 1/s)$ | 2 |
| $s_Z$ | Surface impact on food consumption of host per ectosymbiont attached | $(0, 1)$ | 0.2 |
| $s_{YZ}$ | Surface impact on food consumption of ectosymbiont | $(0, 1)$ | 0.5, 0.6 |
| $\beta_Y$ | Production coefficient of symbiont metabolite by free-living symbiont | $\mathbb{R}_+$ | 2 |
| $\beta_Z$ | Production coefficient of symbiont metabolite by the consortium | $\mathbb{R}_+$ | 0.2, 4 |
| $\delta_U$ | Decay coefficient of host metabolite | $\mathbb{R}_+$ | 1 |
| $\delta_V$ | Decay coefficient of symbiont metabolite | $\mathbb{R}_+$ | 1 |

According to the fixed-point analysis of the two-species resident system, either the host *X* survives alone (as in the previous one-species system) or the host *X* and the symbiont *Y* (both free-living) coexist. Unlike the host species, the symbiont cannot survive entirely on its own (i.e., free of *X*) as it depends on the host metabolite *U* for food. Note that this coexistence is not ectosymbiotic as there is no physical attachment between the host and the symbiont yet. The existence of the interior fixed point of the two-species system depends on how the host’s life history parameters are within bounds governed by the symbiont’s life history parameters (see Appendix A2). The trivial and one-species fixed points are always unstable; while the interior fixed point, if it exists, is always stable unconditionally (Figure 3; for details, see Appendix A2). Apparently, the one-species equilibrium that was previously stable in the one-species system is no longer stable in the presence of a syntrophic species. The syntrophic pair dominates over the host-only system. Hence, metabolite-induced self-inhibition coupled with unidirectional syntrophy can lead to the ecological coexistence of the two involved species (also see Boza et al. (2023)).

**Figure 3.**
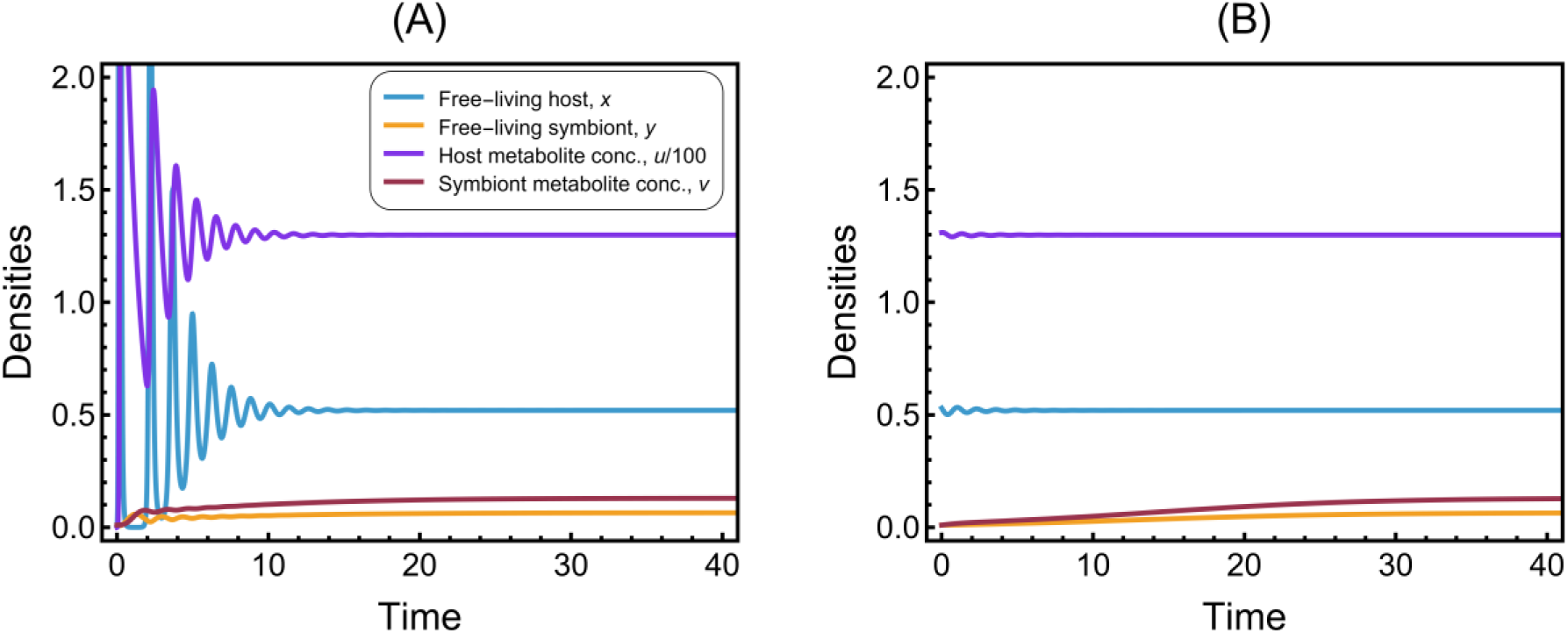
Host and symbiont can stably coexist if symbiont depends on host’s product. (A) Solution of the two-species resident system showing the stability of the unique interior fixed point (*x*^+^, *y*^+^, *u*^+^, *v*^+^) = (0.52, 0.065, 130, 0.13), where both X and Y survive. For an arbitrary point in the neighborhood of the trivial fixed point (0,0,0,0) as the initial condition, the dynamics stabilizes at (*x*^+^, *y*^+^, *u*^+^, *v*^+^) while (0,0,0,0) is unstable. (B) For an arbitrary point in the neighborhood of the one-species resident equilibrium (*x*^∗^, 0, *u*^∗^, 0) as the initial condition, the dynamics again lead to (*x*^+^, *y*^+^, *u*^+^, *v*^+^) while (*x*^∗^, 0, *u*^∗^, 0) is unstable. Arbitrary values of the parameters are as listed in Table 1. See Appendix A2 for more details.

The resident-mutant system with the mutant host can have five ecologically feasible fixed points. The trivial fixed point and the one-species fixed point *E*_1_ = (*x*^∗^, 0,0, *u*^∗^, 0) are always unstable. The two-species resident fixed point *E*_2_ = (*x*^+^, *y*^+^, 0, *u*^+^, *v*^+^) is stable if and only if 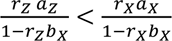, where *r_i_* correspond to the growth rates of the consortial (*Z*) and free-living (*X*) hosts, and *a*_*i*_ and *b*_*i*_ are the growth inhibition factors (see Model and Methods and Appendix A3). The mutant-only fixed point *E*_*M*_ = (0,0, *z̄*, *ū*, *v̄*) exists if and only if the mutant growth rate is less than the maximal growth inhibition and is stable if and only if 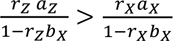. A fixed point *E*_P_ = (0, *ŷ*, *ẑ*, *û*, *v̂*) with symbiont species in free-living and ectosymbiotic forms could also exist conditionally and be stable. There exists no fixed point such that both resident and mutant phenotypes of the host coexist. Figure 4 gives the solutions of the species densities corresponding to the stability of the three possible fixed points. For more details on the existence and stability conditions of the fixed points, see Appendix A3.

**Figure 4.**
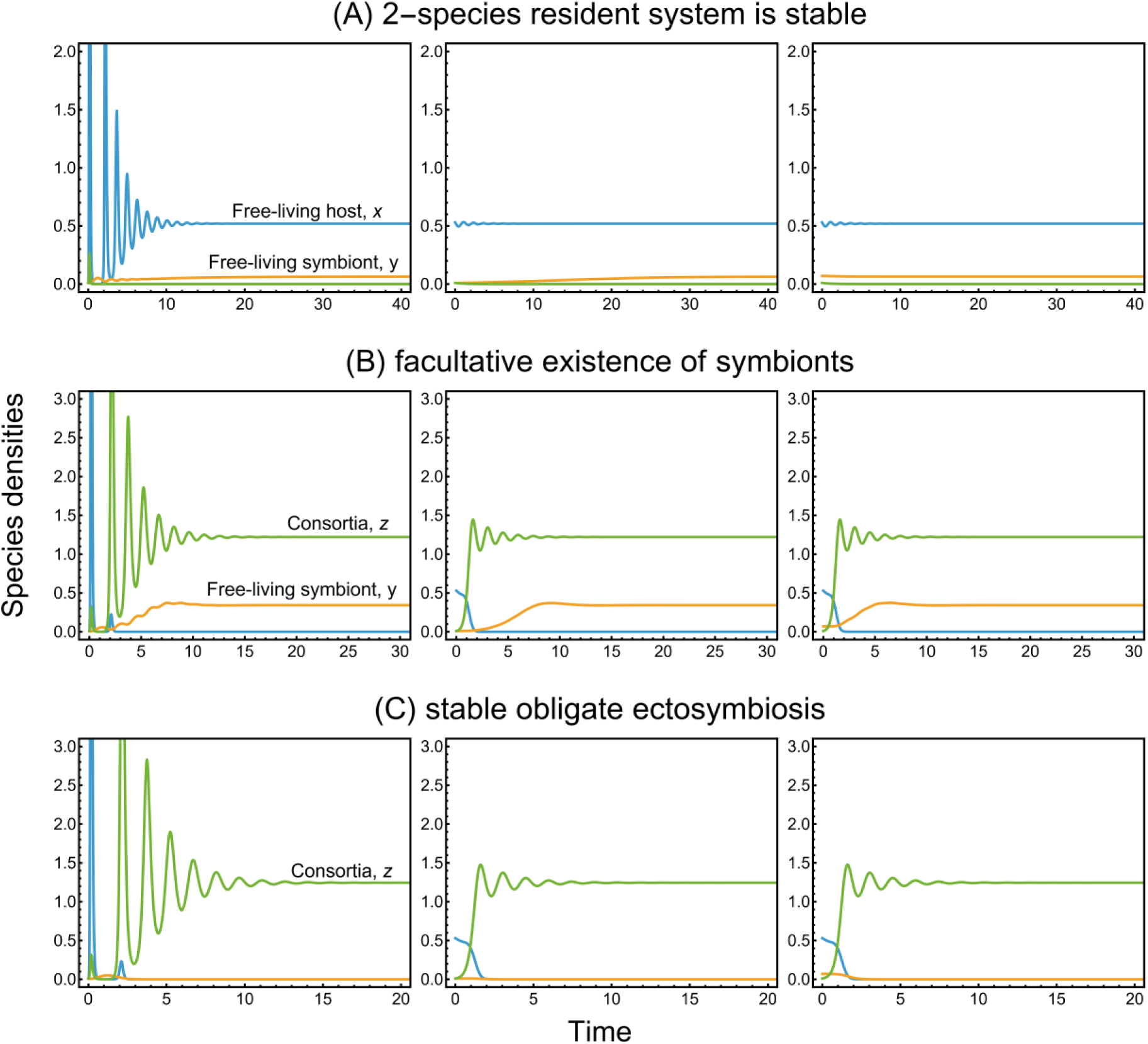
Solution plots corresponding to the species densities in the resident-mutant system dynamics. The arbitrary values of the parameters are listed in Table 1. The initial conditions are arbitrary points in the neighborhood of the trivial, one-species resident and two-species resident equilibria, respectively (column-wise). (A) The dynamics stabilizes at *E*_2_ = (*x*^+^, *y*^+^, 0, *u*^+^, *v*^+^) = (0.52, 0.065, 0, 130, 0.13) for *β*_*Z*_ = 4, *s* = 0.368 and *k*_*YZ*_ = 0.05. (B) The dynamics stabilizes at *E*_*P*_ = (0, *ŷ*, *ẑ*, *û*, *v̂*) = (0, 0.35, 1.23, 165, 0.95) for *β*_*Z*_ = 0.2, *s* = 0.4, and *k*_*YZ*_ = 0.05. (C) The dynamics stabilizes at *E*_*M*_ = (0,0, *z̄*, *ū*, *v̄*) = (0, 0, 1.25, 165, 5) for *β*_*Z*_ = 4, *s* = 0.4, and *k*_*YZ*_ = 0.06. The corresponding metabolite concentrations are plotted separately in Figure 5. See Appendix A3 for more details on stability analysis. Increased occupied surface stabilizes consortia density, and the effective syntrophy parameters of the ectosymbionts promote obligacy.

### Evolution of consortium

The condition for a viable consortium (i.e., mutant invasion condition) is 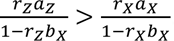. If the invasion condition is not satisfied, the mutant host cannot invade, and the two-species resident equilibrium remains stable; the resident syntrophic system is thus resistant to invasion by the mutant host (i.e., coexistence of free-living species is stable against ectosymbiosis). The mutant invasion condition is satisfied for high values of *a*_*Z*_ relative to *a*_*X*_, where high growth inhibition factor (*a*) implies reduced growth inhibition of the consortia compared to that of the resident hosts. In other words, reduced growth inhibition (or high values of *a*) can facilitate the fixation of consortia as represented in Figure 6B. Now recall that the higher the values of contact surface per ectosymbiont (*s*) and ectosymbiont count (*n*), the higher is the total surface area occupied by the ectosymbionts. As a result, reduced exposure to toxic metabolites or equivalently reduced growth inhibition is realized when the total surface area of contact is high. Figure 6C shows the parameter space in terms of *s* and *n* where the consortia can stabilize. In essence, a mutation in the host that reduces the net growth inhibition by sufficiently protecting it from the self-inhibitory effects of its own metabolites can fixate if these benefits outweigh the costs suffered in the form of a reduced growth rate (Figure 6A). Reduced growth inhibition in the hosts due to the occupied surface area while in ectosymbiosis can be a driving factor in the fixation of the intimate symbiotic consortia, despite the immediate reduction of energetically and metabolically active cell membrane surface. The significance of the area of contact surface in reducing growth inhibition is a reasonable incentive behind the evolution of the ectosymbiotic consortium. Furthermore, this also points to the evolution of enhanced invagination as a mechanism to increase contact surface. Increasing the contact surface likely precedes the eventual engulfment of the syntrophic ectosymbionts that can potentially lead to endosymbiosis, despite entailing a necessary forfeit of useful bioenergetic surface area as species growth rate depends on active (unoccupied) cell surface. Note that the idea generally applies to any microbial syntrophy, not just for the presumed syntrophic origin of mitochondria. Accordingly, we predict that microbial syntrophic partnerships (when the above condition of invasion is met) evolve to increase surface contact area (and also time spent attached to host) through modulating cell-shape, invagination, engulfment, or other means (also see Martin and Müller (1998)). Recently discovered syntrophic Asgards of complex-morphologies (Imachi et al., 2020) suggest that their cellular protrusions may serve exactly this purpose. Our idea is potentially testable in vitro with naturally occurring or artificially assembled symbioses.

### Evolution of obligate ectosymbiosis

Based on the second condition for the stability (exclusively) of the mutant-only equilibrium *E*_*M*_ in Appendix A3, the lower the release of the metabolite *U* into the environment by the consortia, the higher the chance of consortial fixation (i.e., low values of 𝜅 promotes the stability of *E*_*M*_). A high rate of host metabolite consumption by the ectosymbionts in the consortium (*k*_*YZ*_) and a high ectosymbiont count (*n*) reduce the release of metabolite *U* into the habitat and favor the stability of the mutant-only equilibrium (by lowering 𝜅 = (1 − *nk*_*YZ*_)(1 − *ns*_*Z*_)). Similarly, a higher production rate (*β*_*Z*_) of the metabolite *V* by the consortia can also facilitate the stability of *E*_*M*_. In other words, if the ectosymbionts are metabolically active and effectively consume and metabolize the metabolite *U* produced by the consortial host, the mutant-only equilibrium is stable. In this case, the free-living symbiont density collapses due to the constrained availability of resource *U* in the environment and consequent competitive exclusion; hence, fixed point *E*_*P*_ with the facultative existence of the symbiont species (i.e., free-living and ectosymbiotic forms) is not viable. Additionally, in the mutant-only equilibrium, the symbionts are only present in their ectosymbiotic form; the free-living ones go extinct. This means that the stability of the steady state *E*_*M*_ also ensures (and forces) a single lifestyle for the symbionts, thereby developing an obligate ectosymbiosis with the host. In essence, effective metabolic utilization of host metabolite (and subsequent release of symbiont metabolite) by the ectosymbionts could facilitate obligate ectosymbiosis between the mutant host and the symbionts.

### Improved metabolic tolerance of consortia

The mutant hosts in the steady state of the system with only the consortia (stable *E*_*M*_) can tolerate higher concentrations of the host metabolite compared to the resident steady state, i.e., *ū* > *u*^+^ in Appendix A3 (Figure 5). The ectosymbionts protect the mutant host from the toxic metabolite in its environment (produced by itself and conspecifics), increasing their metabolic tolerance. Previously, we observed that the costly mutant could invade the two-species syntrophic resident system if the presence of ectosymbionts can sufficiently help them reduce the effect of metabolic self-inhibition. Also, such a host mutation could form obligate consortia if the dependent ectosymbionts are highly metabolically active. The fixation of the obligate ectosymbiotic consortia can be facilitated by a reduced effective metabolic growth inhibition and a syntrophically solid relationship. The ectosymbionts protect the hosts by consuming the host metabolite effectively, reducing exposure to the host metabolite; thereby enhancing the hosts’ tolerance.

**Figure 5.**
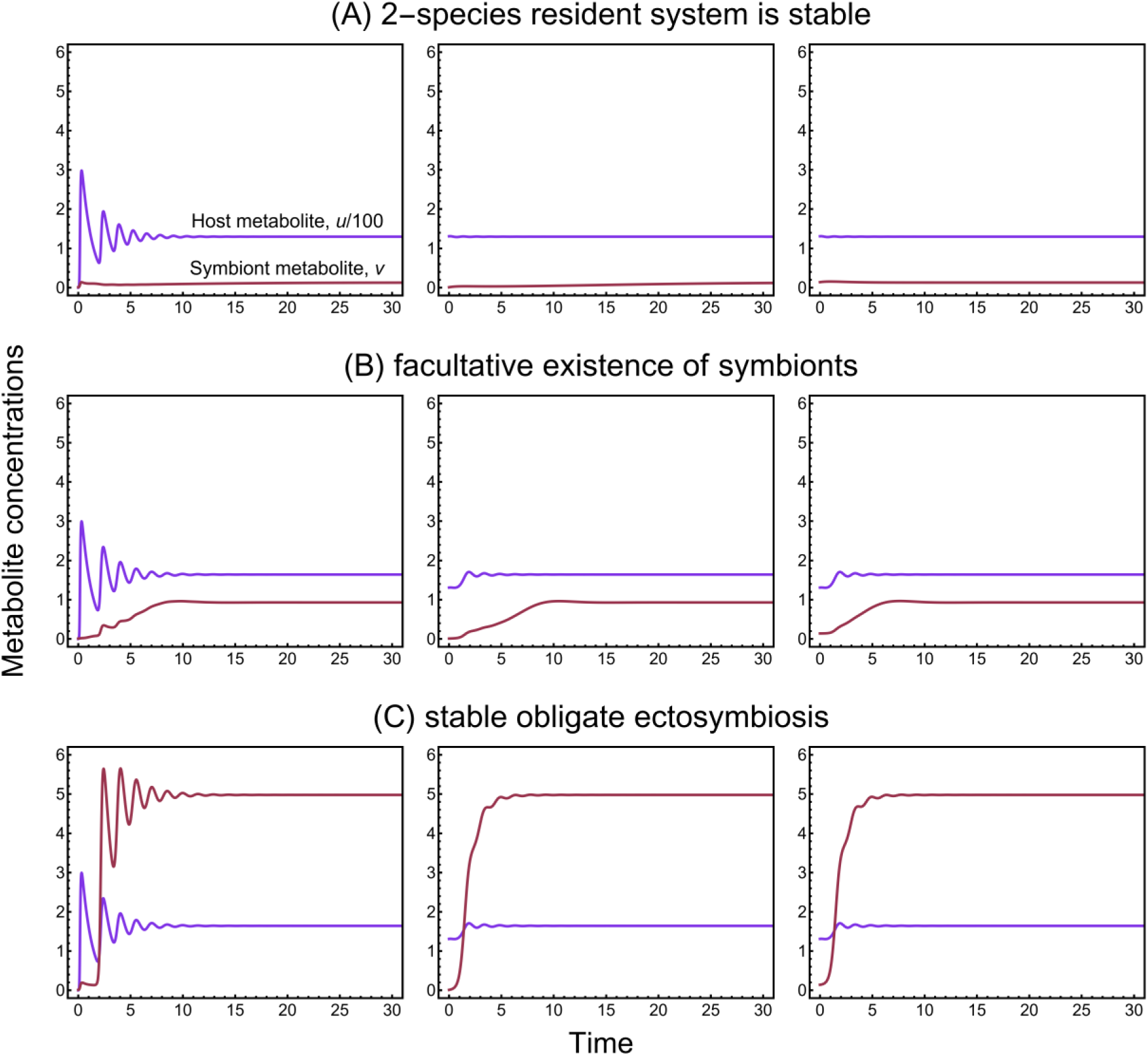
Solution plots corresponding to the metabolite concentrations in the resident-mutant system dynamics. The arbitrary values of the parameters are listed in Table 1. The initial conditions are arbitrary points in the neighborhood of the trivial, one-species resident and two-species resident equilibria, respectively (column-wise). (A) The dynamics stabilizes at *E*_2_ = (*x*^+^, *y*^+^, 0, *u*^+^, *v*^+^) = (0.52, 0.065, 0, 130, 0.13) for *β*_*Z*_ = 4, *s* = 0.368 and *k*_*YZ*_ = 0.05; (B) The dynamics stabilizes at *E*_*P*_ = (0, *ŷ*, *ẑ*, *û*, *v̂*) = (0, 0.35, 1.23, 165, 0.95) for *β*_*Z*_ = 0.2, *s* = 0.4, and *k*_*YZ*_ = 0.05; (C) The dynamics stabilizes at *E*_*M*_ = (0,0, *z̄*, *ū*, *v̄*) = (0, 0, 1.25, 165, 5) for *β*_*Z*_ = 4, *s* = 0.4, and *k*_*YZ*_ = 0.06. The corresponding species densities are plotted separately in Figure 4. See Appendix A3 for more details on stability analysis. The ectosymbiotic consortia can tolerate higher concentrations of the host-metabolite.

**Figure 6.**
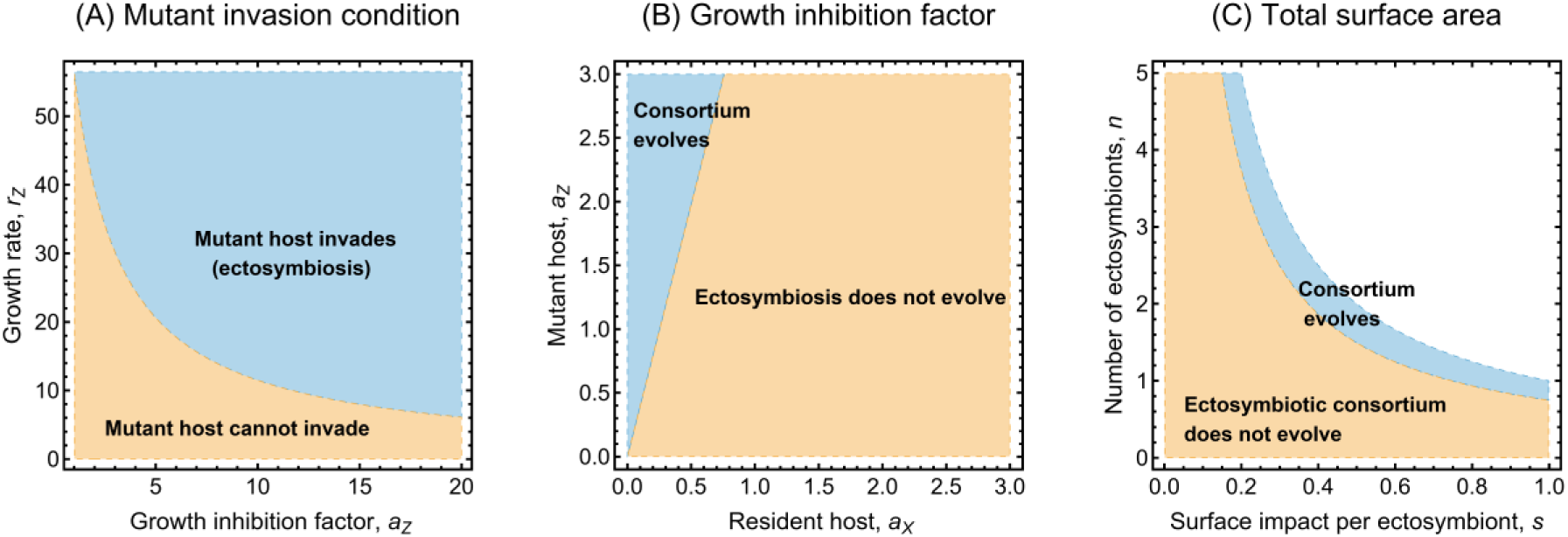
Condition for the fixation of consortia: Region plots representing parameter values corresponding to the stability of resident equilibrium (*x*^+^, *y*^+^, 0, *u*^+^, *v*^+^) and the mutant-only equilibrium (0,0, *z̄*, *ū*, *v̄*). (A) The invasion condition based on the growth rate and growth inhibition factor of the mutant host. (B) Coefficients of growth inhibition in mutant and resident hosts, *a*_*Z*_ against *a*_*X*_. (C) Average ectosymbiont count on a mutant host (*n*) against surface impact on the growth inhibition of the mutant host induced per ectosymbiont on its surface (*s*). For parameters, see Table 1. The ectosymbiotic consortia can potentially fixate on account of the benefit of reduced self-inhibition due to the occupied external surface area of the hosts.

## Discussion

We have modelled the scenario where a host species (a prokaryote, possibly an Asgard archaeon) evolves the means to physically bind individuals of the free-living symbiont species on its plasma membrane to create a consortium. We were interested in whether this interaction can lead to stable coexistence of the consortium-forming host and its symbiont to the point where free-living phenotypes are lost from the population. In our model, the host produces a waste molecule that limits its own growth if accumulating in the vicinity. This molecule, however, is utilized by the symbiont species that can metabolize the inhibitive product; thus, alleviating its effects on the host individual. The syntrophic association is ecologically stable even though the host species was previously viable on its own. The mutant host and the symbiont can form a physically associated, intimate relationship by the host binding the symbiont to its outer surface. The ectosymbiont not only lowers the concentration of the toxic by-product by taking it up but also protects the host from the same inhibitive metabolite produced by the host’s conspecifics in the neighborhood. The magnitude of this effect is the key determinant of the possible outcomes of ecological selection in our model. While this cooperative help has explicit benefits, it also comes with a larger metabolic cost and reduced metabolically active cell surface. The three possible outcomes are as follows: 1) no ectosymbiosis evolves, i.e., the consortium-forming mutant host cannot invade; 2) ectosymbiosis evolves with both free-living and ectosymbiotic symbionts surviving; and 3) only ectosymbiotic consortia survive, resulting in obligate ectosymbiosis. We found that stable ectosymbiotic consortia of mutant host and symbiont can evolve if the net reduction of growth inhibition by the ectosymbionts is large enough to compensate for the inferior growth rate of the mutant hosts compared to the residents.

Our model assumes that the mutant host has a novel mechanism expressed on its external surface to bind symbiotic cells. While in biofilms, associated prokaryotes stay together because of the matrix they produce (Dunne, 2002); there are many dedicated mechanisms for prokaryotic cells to attach to each other. For example, archaea can attach to surfaces or to other cells by hami- (Ng et al., 2008), flagellae- or pili-like structures (Fröls et al., 2008; Näther et al., 2006; Tripepi et al., 2010). In biofilms of *Pyrococcus furiosus* and *Methanopyrus kandleri*, apart from pili, a yet unknown contact molecule acts as binder (Schopf et al., 2008). Bacterial dynamin-like proteins 1 (DLP1) bind to membranes, and the DLP1/DLP2 complex can tether two membranes together (Bramkamp, 2018). Coiled-coil structures with globular heads at their end can connect two membranes at a distance (Chia and Gleeson, 2014) (they are mostly involved in the functioning of the Golgi apparatus). Based on the above biological examples, it is not far-fetched to assume a small change in a surface protein (or a cytosolic protein that begins to be expressed on the membrane) of the host initiates a more intimate connection with a partner cell.

The connection with the host reduces the active cell surface of it, thus the connection must come with a metabolic price. Most prokaryotes feed through their plasma membranes, and their metabolic production is thus directly dependent on surface area. In prokaryotes, especially in autotrophs, growth rate is strongly dependent on cell surface area (Schavemaker and Muñoz-Gómez, 2022). However, we also point out that our model is not specific on molecular surface structures in binding partner cells. As a matter of fact, any surface- mechanism that keeps partner cells attached to the host suffices. For example, membrane invaginations or protrusions can serve to capture the partner cells and keep them attached to the host. While there exist such mechanisms (Imachi et al., 2020; Pande et al., 2015), admittedly, it is unknown if they are capable of ensuring long-term coexistence of parties. Such mechanisms, if they were ever involved in establishing obligate ectosymbioses, are beneficial over simple surface attachments due to at least two reasons. First, they may be able to compensate for the loss of active cell surface for the host; and second, they may better ensure vertical inheritance if the symbiont is enclosed such that it cannot escape. Either way, the model indicates the significance of the area of surface contact between parties in the evolution of ectosymbioses.

Our model ignores the individual dynamics (reproduction, death, escape, etc.) of the attached ectosymbionts. While this is a must for the model to remain tractable, it is also justifiable. We assume that as the host cell reproduces, it doubles its genome and materials. This means that with every division, the cell membrane is also doubled in size. Accordingly, we assume that as the host increases its surface area, the number of attachment points also increases. Immediately after division, a cell surface of *A* can hold *n* ectosymbionts; while immediately before division, we assume that 2*A* can hold 2*n* ectosymbionts. Assuming that the membrane is fluid, attachment points are distributed evenly on the surface; as a result, daughter cells after division receive equal amounts: *n* ectosymbionts. Any excess replication of the ectosymbionts when attached to the surface is ignored as, if offspring cannot fit over the host, it is released to the external symbiont pool which is assumed to be much larger. Admittedly these are simplifications, but this approach does not require synchronized cell cycles of host and symbiont. Note that we do not assume perfect vertical inheritance as we do not specify if the same symbionts are present always or occupy specific attachment points, they may come and leave independently and randomly. The important point is that a number of them are always attached to the host after a random assortment – implementing thus a poor man’s passive vertical inheritance.

In our model, the different host phenotypes (consortial and nonconsortial) do not compete for any resource (neither for their primary food nor for symbionts). Any such competition would further help the consortial host in channeling away free symbionts, reducing external symbiont population. This in turn increases inhibition for the non-consortial hosts that rely explicitly on free-living symbionts for reducing the toxic metabolite concentration. Our results show that there is no mixed coexistence where both host phenotypes can survive.

In a previous work (Krishnan et al., 2024), a theoretical model with a unidirectional syntrophic relationship of two physically independent host and symbiont species was considered in a different selection scenario. Instead of metabolic growth inhibition, the population growth was limited by intraspecies density-dependent competition. Similar to the study in this paper, the possible invasion of a mutant capable of forming an ectosymbiotic consortium was analyzed; however, there it was assumed that the mutation emerges in the symbiont species and not in the host, serving as a crude model of the parasitic invasion theory (Searcy, 2003; White et al., 2018). It was observed that the metabolic activity and the additional costs incurred by the mutant symbiont (compared to the resident) determine the stability of the consortium. However, mutant symbionts, while replacing the residents, were found to survive in both free-living and ectosymbiotic forms. In other words, no obligate association of the symbiotic consortium – from the perspective of the symbionts – was observed in the model (Krishnan et al., 2024). In this work, however, we have shown that an obligate ectosymbiotic association can evolve.

Despite significant advances in research, the origin of the eukaryotic cell still remains an enigma; most importantly, the basis of the permanent association and the mechanism of entry of the bacterial partner remains unknown. A common idea in most theories about the origin is that syntrophy *somehow* led to the endosymbiosis of the early partners. This idea is prevalent in many forms (Baum and Baum, 2014; Imachi et al., 2020; López-García et al., 2017; López-García and Moreira, 2023, 2020; López-García and Moreira, 1999; Martin et al., 2015; Martin and Müller, 1998; Searcy, 2003; Spang et al., 2015), despite the fact that no syntrophic prokaryotes are known yet that have concluded their merger by provably forming a higher level unit of evolution. Most of the known endosymbionts of unicells are known to have entered their host via the phagocytotic mechanism, e.g. all plastids (Bhattacharya et al., 2012; Keeling, 2013, 2010; Zimorski et al., 2014). However, prokaryotes lack a homologous mechanism which may account for the lack of endosymbioses among them. While there is at least one known analogous endocytic mechanism in prokaryotes (Shiratori et al., 2019), it is neither on par with eukaryotic phagotrophy nor is it known to lead to (or facilitate) endosymbiotic integration, and – most importantly – there are no such mechanisms known in Asgards, the closest archaeal relatives of eukaryotes (Spang et al., 2019). While the last eukaryotic common ancestor might have already had phagocytotic capabilities (Bowles et al., 2023; Gawryluk et al., 2019), we do not yet know if it evolved before or after the integration of the mitochondrial ancestor (also see Richards et al. (2024), Bremer et al. (2022), and Mills (2020)). Some Asgards, though possess rudimentary cytoskeleton and relevant proteins (Charles-Orszag et al., 2024; Imachi et al., 2020; Rodrigues-Oliveira et al., 2023; Wollweber et al., 2025), have not shown endocytic capabilities so far (Mills, 2020). Consequently, as of now, we still don’t know the mechanism of entry of the bacterial symbiont into the ancestor of eukaryotes. The phagotrophic scenario (Cavalier-Smith, 2010, 2002; Cavalier-Smith and Chao, 2020) was already explored in a model by Zachar et al. (2018). If, however, we assume phagotrophy was a late eukaryotic invention after mitochondria (e.g., Martin et al. (2017)), then we must account for an alternative mechanism on how the host has captured and internalized its symbiont. Furthermore, we must point out that metabolic dependence, capture (or binding) and internalization are distinct steps that the different scenarios explain in different orders.

Besides phagotrophy, the two main contenders of endosymbiotic inclusion are parasitic invasion (White et al., 2018) and syntrophic engulfment (Baum and Spang, 2023; Baum and Baum, 2014; Imachi et al., 2020). While we note that *engulfment* is not a very well-defined term, there are nevertheless at least two reasons why syntropy seems to be the more likely candidate for initial interaction of ancestral parties. First, parasitic infection is a disadvantage to the host that is unlikely to be selected for; contrarily, a neutral or even beneficial metabolic interaction seems to be a better starting point, especially if it is already mutual at the start (López-García and Moreira, 1999; Martin and Müller, 1998; Moreira and López-García, 1998; Sousa et al., 2016; Spang et al., 2019). Second, metabolic syntrophy is ubiquitous among prokaryotes to the point where it is intriguing why it has never triggered more endosymbioses that we know of (Zachar and Boza, 2022, 2020). Here, we focused on the latter theory, exploring the possibility that if metabolic dependence, at least unilateral, has already evolved between the parties, then physical attachment can lead to obligate partnership. We have tested the hypothesis that ectosymbiosis can evolve from metabolic cooperation.

A particularly relevant hypothesis seems to be the inside-out origin of eukaryotes (Baum and Spang, 2023; Baum and Baum, 2014), also consistent with the entangle-engulf-endogenize hypothesis (Imachi et al., 2020). This scenario assumes an initially metabolic interaction, likely mutually dependent and beneficial for both parties, that evolved to more intimate physical entanglement. Ultimate integration happened when the host’s protrusions evaginated and enclosed the symbiont(s) and ultimately fused with itself, endogenizing the captured partners. There is no proof yet that such a fusing mechanism happens or could work in prokaryotes; the basic idea is the same as in all syntrophic scenarios: *somehow*, one partner was endogenized without an explicit phagocytotic mechanism. Without trying to explain the last bit, here we focused on how the metabolic interaction of two free-living species can trigger obligate membrane-bound ectosymbiosis.

Our model explicitly assumes that genetic dependence has already evolved in the sense that symbionts depend on host cell products (however, hosts need not depend on symbiont as it can live freely, though with increased self-inhibition). These assumptions are well-supported by real-world examples. Association between prokaryotes, particularly between Archaea and Bacteria is ubiquitous (Wrede et al., 2012). The association can often be syntrophic, where the inhibitive waste product of one species is consumed by another (Bull and Harcombe, 2009; Estrela et al., 2012; Fennell and Gossett, 1998; Kreikenbohm and Bohl, 1986; Oliveira et al., 2014). For example, photosynthetic *Chlorobi* bacteria attach to a larger *β*-proteobacterium to form epibiotic consortia (Cerqueda-García et al., 2014; Liu et al., 2013; Overmann and van Gemerden, 2000), or filamentous, sulphur-oxidizing γ-proteobacteria of the genus *Thiothrix* associates with coccoid euryarchaeota forming pearl-like structures (Moissl et al., 2002; Rudolph et al., 2004, 2001). In the latter, the bacterial partner possibly gives metabolites to the archaea and shields them from the aerobic environment. In this system, it is the bacteria which shields a multitude of smaller archaea. However, small, sulfur-oxidizing γ-proteobacterial cells can also be seen covering the surface of the giant archaeal cells of *Giganthauma karukerense* (Muller et al., 2010). This arrangement is morphologically identical to the one proposed in our model. Even more similar examples are discovered among syntrophic Asgards too (Farag et al., 2020; Hager et al., 2025; Imachi et al., 2020; Spang et al., 2019).

A special type of cross-feeding is the detoxification of inhibitory molecules, and the previous examples hint not only about potential metabolic interactions (syntrophy) and their benefits, but also about the adverse effect of products on one partner that can be eliminated by the other. For example, *Escherichia coli* produces acetate as a by-product of metabolizing glucose, which by changing the anionic environment, hinders its growth (Pinhal et al., 2019). Another example, yeast (*Saccharomyces cerevisiae*) produces alcohol as a by-product of breaking down glucose, yet excess concentration of alcohol becomes toxic, inducing severe stress for the yeast (Ding et al., 2009). *Bacillus cereus* MLY1 strain can degrade the polluting compound tetrahydrofuran; however, the resulting acidic environment makes it very inefficient. In the presence of *Rhodococcus ruber* YYL, metabolic cross-feeding, especially the removal of acidic compounds by *R. ruber*, makes the co-culture more efficient at biodegrading tetrahydrofuran (Liu et al., 2019).

The fully obligate ectosymbiosis (an outcome of our model) enables and may ultimately lead to vertical inheritance of the symbiont, thus creating a higher-level unit of selection (this transition remains to be seen either in the lab or in simulations). Moreover, successful host-symbiont integration also depends on the timing of evolution of the relevant innovations, for instance, the synchronization in replication, perfect (or effective) vertical inheritance and the dependence between the hosts and symbionts in consortium (Athreya et al., 2025). With the fixation of the new evolutionary unit, selection for better-integrated interactions and novel features can become achievable. An upcoming paper is taking the modelling a step further towards physical inclusion, discussing the possibility of improved contact surface (as in invagination) as a mechanism to counter growth inhibition.

## Model and Methods

### Malthusian growth rate

Since the model formulation relies on metabolic consumption and the population’s intrinsic growth is metabolite-dependent, we utilize the Malthusian growth definition explained in Krishnan et al. (2024), based on deterministic and uniformly continuous resource consumption and cell replication of an asexually reproducing unicellular organism, as:

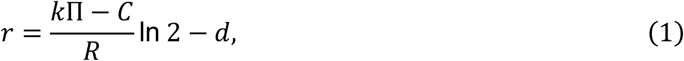

where *k* is the rate of resource consumption, Π the available resource concentration, *C* the cost of living (per capita consumption required to sustain cell), and *d* the intrinsic death rate. The organism is assumed to replicate (e.g., binary splitting (Gao et al., 2022)) upon acquiring a threshold amount of biomass represented as cost of reproduction, *R*.

The organism, while benefiting from consuming the available resource, in turn produces a metabolic waste. The concentration of the metabolic product of the organism negatively impacts its growth, i.e., excess concentration of the by-product is toxic to the producer (e.g., alcohol produced by yeast (Ding et al., 2009)). Accordingly, we model the net growth rate of the producer declining with rising concentration of the produced metabolite (growth inhibition), bounding growth and potentially stabilizing the population.

### One-species resident system: self-inhibition by metabolite production

First, we analyze the behavior of a single-species resident system to examine whether growth inhibition stabilizes the producers’ population as expected (Figure 1A). We assume species *X* (would-be host) acquires their primary resource, 𝛹 from a replenishing source in the environment having an available constant concentration 𝜓. Based on the description of the Malthusian growth rate, 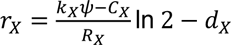 is the per capita growth rate of species *X*. The organism then produces a metabolic product *U* and releases it into the external environment as waste at a rate proportional to its food consumption. The cumulative metabolic product in the environment is toxic to the organism and limits its growth according to a non-linear function motivated by the Monod equation (Monod, 1949):

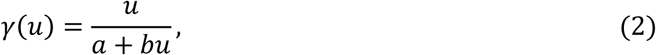

where *u* is the variable concentration of the released product, *b* defines the value of maximal inhibition, and 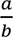 is the half-velocity constant. We vary these factors to differentiate the growth inhibition for different species. We also assume that the metabolite *U* decays at a constant rate (𝛿_*U*_). We propose the following deterministic model for the dynamics of a single host species *X* (with population density *x*) and metabolite *U* using a set of non-linear differential equations:

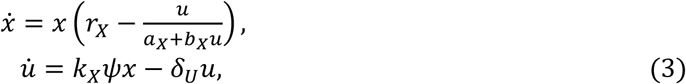

where all the parameters are strictly positive.

### Two-species resident system: syntrophy evolves

Now consider a second species *Y* (symbiont), capable of forming a unidirectional metabolic relationship with species *X* (host) (Figure 1B). The symbiont *Y* consumes the metabolic product *U* of *X*. We assume that the host metabolite *U* released into the habitat is diluted in the medium (having volume Ω); hence, 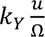 is the effective per capita consumption of metabolite *U* in the medium by the free-living symbionts. 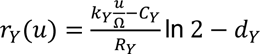 is the metabolite-dependent per capita growth rate of species *Y*. The symbiont, in turn, releases waste *V* into the environment at a constant rate (*β*_*Y*_). Just as *U* inhibits *X*, *V* is toxic to the free-living symbiont and inhibits its growth. *V* also decays at a constant rate 𝛿_*V*_. Denote *y* as the population density of species *Y*, and *v* as the concentration of the metabolite *V* in the habitat. The two-species extended system of the syntrophic pair is as follows:

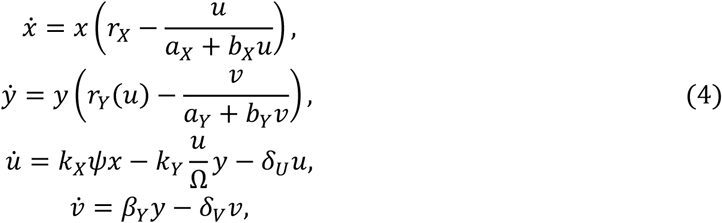

where all the parameters are strictly positive. The resource-consumer dynamics here assumes no timescale separation between the species’ population densities and the metabolite concentration dynamics (also see Boza et al. (2023)). The syntrophic resident (monomorphic) system is analyzed to check for the stable coexistence of the host-symbiont pair.

### Mutant host invasion: ectosymbiosis evolves

Next, we consider an invasion by a mutant phenotype of the host into the two-species syntrophic resident system (Figure 1C). Mutant invasions are usually rare, and consequently, we assume that the mutant emerges at a stable state of the resident system (Cressman and Garay, 2003a, 2003b), and back mutation is unlikely. As a result, it is reasonable to assume that all the mutant hosts could have descended from a single mutant individual. The mutant hosts (denoted *Z*) have novel structures that enable them to bind the free-living symbionts *Y* on their external surface and initiate an ectosymbiotic association. We consider that the free- iving symbiont pool is considerably larger than the host population. The mutant hosts are assumed to always form (and exist only in) the consortia; hence, the two terms are interchangeable in our context. This mutant host along with the ectosymbionts (*consortium*) is considered a higher-level organization (Maynard Smith and Szathmáry, 1995; Rainey and Kerr, 2010; Rainey and Rainey, 2003; Szathmáry, 2015). The consortium replicates based on the host characteristics, and its offspring remain in the same population group owing to a random assortment of symbionts on the host surface before and after replication. For the sake of simplicity, we do not explicitly model the ectosymbiont dynamics, particularly interspecific interactions leading to their formation, replication and dissociation (more explanation in Discussion).

The mutant hosts also consume resource Ψ and produce the toxic waste metabolite *U*, analogous to the resident hosts. The mutation to associate symbionts to the plasma membrane is, however, assumed to be costly, i.e., the costs of living and reproduction of the consortia *Z* are higher than that of the resident *X* (*C*_*X*_ < *C*_*Z*_ and *R*_*X*_ < *R*_*Z*_). Additionally, accounting for the effect of the symbiont metabolite on the consortium’s net growth rate, we consider a higher intrinsic death rate for the consortium, i.e., *d*_*X*_ < *d*_*Z*_ to indirectly reflect that the consortium is worse off in that aspect. Otherwise, we assume the mutant host is identical to the resident species. The presence of symbionts on the surface benefits the mutant host by reducing the concentration of toxic metabolites released by the *X* and *Z*. However, the presence of the ectosymbionts also constrain the taking up (consumption) of resource Ψ by *Z*. Thus, the consumption rate of the mutant host is lower than that of the resident host (*k*_*X*_ > *k*_*Z*_). The rationale behind these two effects is detailed below.

#### Reduced surface reduces growth inhibition

The attached ectosymbionts decrease the free external surface area of their mutant host. As a result, there is an effective reduction in the external surface exposed to the metabolite *U* in the environment. While in ectosymbiosis, contact decreases the growth inhibition of the mutant hosts induced by concentration of *U* in the environment. Let us denote by *s* ∈ (0,1) the surface area of contact (or a measure of impact on the growth inhibition of the host) per ectosymbiotic cell attached and the ectosymbiont count by *n* ∈ [1, 1/*s*). Without loss of generality, we consider unity as the maximum of total surface area of contact, 0 < *s* < *ns* < 1, where *ns* correspond to the surface of the mutant host covered by the total *n* attached symbionts. The mutant host thus has an advantage over the free-living residents in terms of its reduced exposure to the toxic metabolite, which can be represented as follows:

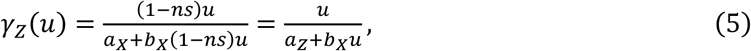

where 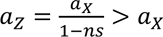 and it increases with increasing total contact area. Recall that, for free-living resident hosts, 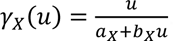

### Reduced surface reduces food consumption

The influx of the metabolites into the cell body is also constrained in proportion to the reduction in the free surface of the mutant host. Thus, the host’s food reaches the interior of the mutant host with ectosymbionts at a lower rate. The consumption of the mutant host in the consortium is lowered by the presence of ectosymbionts. Let *s*_*Z*_ ∈ (0,1) denote the measure of reduction in the consumption rate of Ψ induced by a single ectosymbiont. The consumption rate of the mutant host with *n* attached symbionts can then be given by:

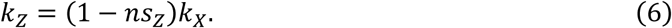

Larger contact surface corresponds to higher survival of the mutant host by protecting it from the external toxic metabolite but simultaneously corresponds to lower fecundity by constraining food consumption. Thus, there is a trade-off between growth inhibition and food consumption of the mutant hosts. Consequently, the integration of the effects of occupied surface area on these two traits determines the mutant host’s net benefit, while incurring additional costs of living and reproduction compared to the resident host.

Similarly, an ectosymbiont could consume the resource *U* either from the habitat through its free surface (as *U* dissolved in the environment) or directly from its mutant host without any dilution, i.e., total per capita consumption of an ectosymbiont is 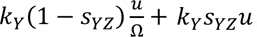, where *s*_*YZ*_ is the surface impact on the consumption of ectosymbionts (subscript *YZ* correspond to the ectosymbionts, i.e., species *Y* involved in consortia *Z*). For the free-living symbionts, entire surface is free, and the expression reduces to 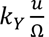. For simplicity, we assume that for each attached symbiont, the consumption of the metabolite directly from its host is much larger than that of the diluted metabolite, i.e., 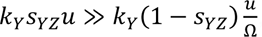 and consequently, we ignore the attached symbiont’s consumption from the habitat. Denoting *k*_*YZ*_ as the consumption rate of the ectosymbionts, *k*_*YZ*_ = *k*_*Y*_*s*_*YZ*_ < *k*_*Y*_, i.e., the consumption rate of the ectosymbionts is lower than the consumption of the free-living symbionts due to reduced free surface. Moreover, the ectosymbiont consumption rate is proportional to the surface area of contact. We assume that the mutant host in consortium can produce more quantity of metabolite *U* than the maximum collective consumption capability of *n* ectosymbionts, i.e., *nk*_*YZ*_ < 1. The remaining metabolite after direct consumption by the ectosymbionts is released into the medium by the consortia at the rate (1 − *nk*_*YZ*_)*k*_*Z*_𝜓 = (1 − *nk*_*YZ*_)(1 − *ns*_*Z*_)*k*_*X*_𝜓. Recall the concentration of metabolite *U* produced by the consortium is proportional to its food consumption. For simplicity, let 𝜅: = (1 − *nk*_*YZ*_)(1 − *ns*_*Z*_). The consortia also release the metabolite *V* into the medium at a constant rate *β*_*Z*_. The effect of the metabolite *V* on the ectosymbionts and the consortia is collectively incorporated in the consortia’s reduced growth rate indirectly. We assume that the symbiont metabolite does not have a direct impact on the consortium, i.e., the metabolite-dependent growth inhibition of the consortium is based on the host metabolite only.

### Resident-mutant system: fixation of consortia

Based on the model descriptions, the mutant host that forms the consortium has an advantage over the resident in terms of reduced exposure to *U*. However, the mutant phenotype suffers additional costs due to constrained food consumption and higher costs of living and reproduction, which are reflected in a reduced growth rate. Now using the dynamical approach for population dynamics as in the resident system, we analyze whether this mutant host can invade the two-species, syntrophic resident system and replace the resident host phenotype (Cressman et al., 2020). The mutant host (population density, *z*) is introduced as a new dimension to the 4D resident system and the extended system is as follows:

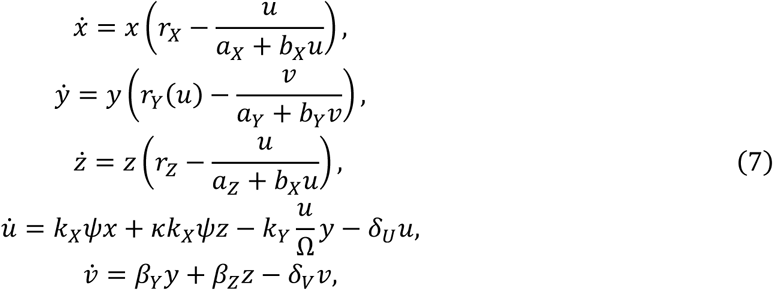

where all parameters are strictly positive.

### Evolutionary substitution: a dynamical approach

The model analysis is based on integrating the influence of density-dependent, multi-species ecological dynamics and environment on evolutionary processes (Cressman and Garay, 2003a, 2003b). Most of the assumptions of the model are analogous to the adaptive dynamics approach (Dieckmann and Law, 1996; Geritz et al., 1998, 1997); for instance, the timescale separation between the ecological dynamics and occurrence of rare mutation. However, metabolite-dependent population growth and growth inhibition is effectively modelled using a resource-consumer-based dynamics (Boza et al., 2023; Chesson, 1990; Holland et al., 2005; MacArthur, 1970). Also, our definition of the mutant host phenotype and its invasion motivated us to use a much flexible coevolutionary dynamical approach (Cressman, 1992; Cressman et al., 2020; Cressman and Garay, 2003a, 2003b; Garay, 2007). The resident-mutant coevolutionary dynamics is built based on the resident ecological dynamics; introducing mutant dynamics as a new dimension to resident systems creates extended ecological dynamics, where the mutant invasion and fixation is based on the outcomes of ecological selection. Linear stability analysis of the steady states (fixed points) of the resident-mutant dynamics points to the potential states to which the system can evolve. *Evolutionary substitution* corresponds to the stability of the steady state where the mutant survives while replacing the respective resident; the costly mutant dominates the resident as the resident-only steady state is simultaneously unstable (Cressman et al., 2020). The substitution of the resident by the invading, consortia-forming mutant translates to the fixation of the consortia. While analyzing the dynamics of the systems, we are only interested in the stability of the steady states. Even though the system could have complicated dynamics (for instance, limit cycles), this exceeds the scope of our work as it would require varying resident densities when the mutant is introduced. This is clearly a simplification, and we prefer to first focus only on the case where the mutant is introduced to a stable resident system with unique steady states. Assuming the mutant to be rare in density, analysis of the local behavior rather than the global behavior can also be justified. It is indeed worthwhile to analyze the system’s behavior in entirety.

## Acknowledgements

This project has received funding from the European Union’s Horizon 2020 Research and Innovation Programme under the Marie Skłodowska-Curie grant agreement number 955708 (NK, ÁK, CSG, JG). IZ acknowledges funding from MTA Bolyai János Research Scholarship #BO/00570/22/8 and John Templeton Foundation grant ID 63451 “Direction, agency and function in the evolution of symbiotic integration”. NK also acknowledges helpful discussions with Prof. Mark Broom.

## Author Contributions

**NK**: Conceptualization, Methodology, Visualization, Formal analysis, Writing – original draft and Writing – review & editing

**IZ**: Conceptualization, Methodology, Visualization, Writing – original draft and Writing – review & editing

**ÁK**: Conceptualization, Methodology, Writing – original draft and Writing – review & editing

**CSG**: Conceptualization, Methodology and Writing – review & editing

**JG**: Conceptualization, Methodology, Supervision, Writing – original draft and Writing – review & editing

## Appendix

## Appendix A1: Stability analysis of one-species system

The one-species (host; species *X*) system dynamics is defined as follows:

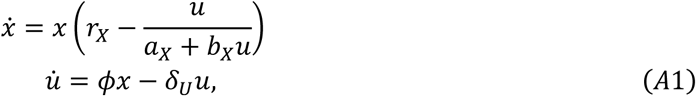

where 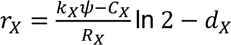 and 𝜙 = *k*_X_ 𝜓. The system can have two biologically feasible fixed points: trivial (0, 0) and a unique interior fixed point 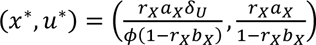 that exists if and only if 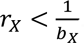. The Jacobian matrix corresponding to the system for arbitrary (*x*, *u*) is

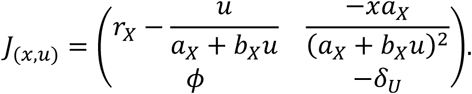

*J*_(0,0)_ has a positive eigenvalue; hence, (0, 0) is unstable. The real part of both eigenvalues of *J*_(*x*_∗_,*u*_∗_)_ is always negative; hence, (*x*^∗^, *u*^∗^) is always stable, if it exists.

## Appendix A2: Stability analysis of two-species system

The two-species (host-symbiont syntrophic pair; species *X* and *Y*) system dynamics is defined as follows:

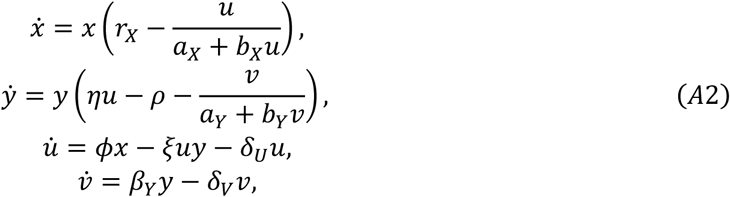

where 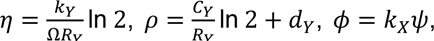 and 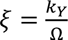 (rest of the parameters are as in Appendix A1). The system can have three biologically feasible fixed points: trivial (0, 0, 0, 0), one-species fixed point (*x*^∗^, 0, *u*^∗^, 0), and an interior fixed point (*x*^+^, *y*^+^, *u*^+^, *v*^+^) where

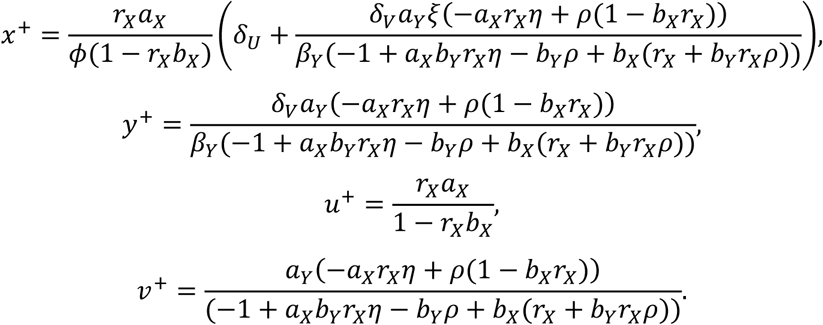

The fixed point (*x*^+^, *y*^+^, *u*^+^, *v*^+^) exists if and only if 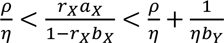 and 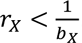. The Jacobian matrix corresponding to the system for arbitrary (*x*, *y*, *u*, *v*) is

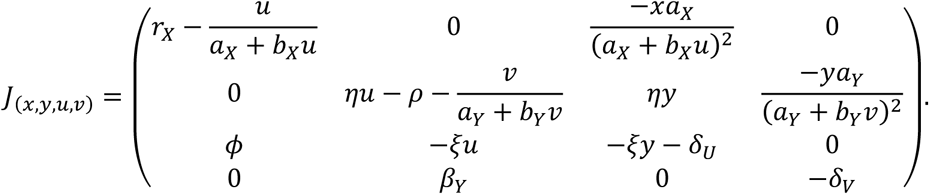

*J*_(0,0,0,0)_ and *J*_(*x*_∗_,0,*u*_∗_,0)_ have a positive eigenvalue as *r*_*X*_ > 0 and 𝜂*u*^∗^ > 𝜌, respectively; hence, corresponding fixed points are unstable. The Jacobian matrix corresponding to the system at (*x*^+^, *y*^+^, *u*^+^, *v*^+^) is

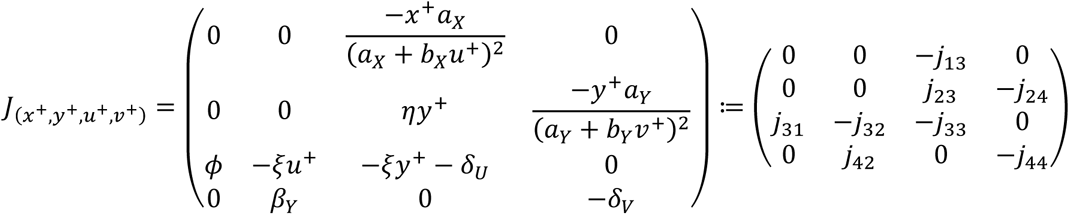

where in the redefined matrix elements of *J*_(*x*+,*y*+,*u*+,*v*+)_, *j*_13_, *j*_23_, *j*_24_, *j*_31_, *j*_32_, *j*_33_, *j*_42_, and *j*_44_ are positive-valued. Now, the characteristic equation of *J*_(*x*+,*y*+,*u*+,*v*+)_, for 𝜆 denoting eigenvalues, can be expressed in the form

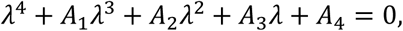

^where *A*^1 ^= *j*^33 ^+ *j*^44^, *A*^2 ^= *j*^13^*j*^31 ^+ *j*^23^*j*^32 ^+ *j*^24^*j*^42 ^+ *j*^33^*j*^44^, *A*^3 ^= *j*^24^*j*^33^*j*^42 ^+ *j*^13^*j*^31^*j*^44 ^+ *j*^23^*j*^32_44_, and *A*_4_ = *j*_13_*j*_24_*j*_31_*j*_42_. Here, as *A*_1_ > 0, *A*_2_ > 0, *A*_3_ > 0, *A*_4_ > 0, *A*_1_*A*_2_ > *A*_3_, and 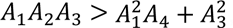 using the Routh-Hurwitz criterion (Edelstein-Keshet, 2005; Gantmacher, 1960), real part of all eigenvalues of *J*_(*x*+,*y*+,*u*+,*v*+)_ is negative; hence, (*x*^+^, *y*^+^, *u*^+^, *v*^+^) is always stable, given it exists.

## Appendix A3: Stability analysis of resident-mutant system

The resident-mutant system dynamics is defined as follows:

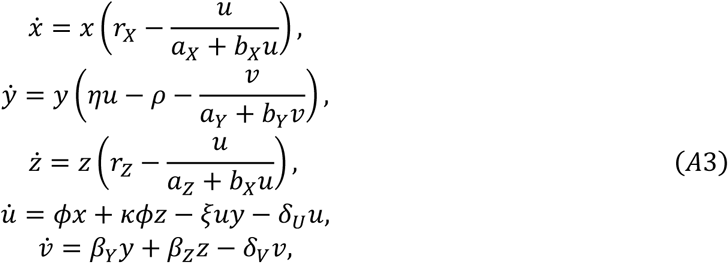

where 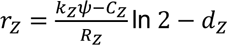 and 𝜅 = (1 − *ns_Z_*)(1 − *nk_YZ_*) (rest of the parameters are as in Appendix A2). The system can have five feasible fixed points: *E*_0_ = (0,0,0,0,0), *E*_1_ = (*x*^∗^, 0,0, *u*^∗^, 0), *E*_2_ = (*x*^+^, *y*^+^, 0, *u*^+^, *v*^+^), *E*_*P*_ = (0, *ŷ*, *ẑ*, *û*, *v̂*) and *E*_*M*_ = (0,0, *z̄*, *ū*, *v̄*). The Jacobian matrix corresponding to the system for arbitrary (*x*, *y*, *z*, *u*, *v*) is ^*J*^_(*x*,*y*,*z*,*u*,*v*)_

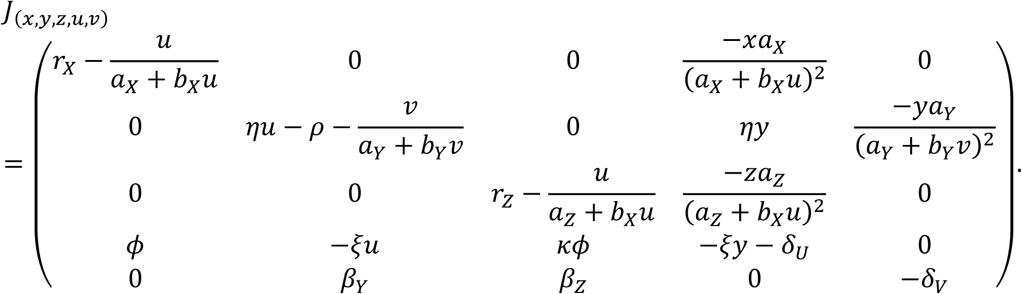

*E*_0_ and *E*_1_ are always unstable. The resident equilibrium *E*_2_ = (*x*^+^, *y*^+^, 0, *u*^+^, *v*^+^) is stable if and only if 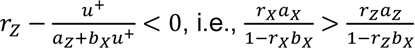. The mutant-only equilibrium 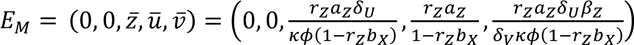 exists if and only if 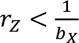 and is stable if and only if

1. 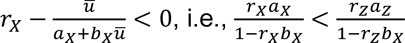 and
2. 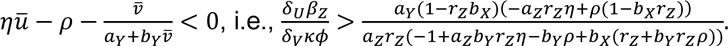

Assuming existence of *E*_2_ and *E*_*M*_, if condition 1 is not satisfied, the resident equilibrium *E*_2_ is stable; and, if only condition 2 is not satisfied, then *E*_*P*_ = (0, *ŷ*, *ẑ*, *û*, *v̂*) can also exist and be stable.

## Notes

### Competing Interest Statement

The authors have declared no competing interest.

### Summary of Updates

The manuscript has been revised with major changes to the Introduction and Discussion; Title is updated; figures are updated; a new author is added.

